# Structural basis for the interaction of SARS-CoV-2 virulence factor nsp1 with Pol α - Primase

**DOI:** 10.1101/2021.06.17.448816

**Authors:** Mairi L. Kilkenny, Charlotte E. Veale, Amir Guppy, Steven W. Hardwick, Dimitri Y. Chirgadze, Neil J. Rzechorzek, Joseph D. Maman, Luca Pellegrini

**Author notes:** Current address: The Francis Crick Institute, London, NW1 1AT, UK.

## Abstract

The molecular mechanisms that drive the infection by the SARS-CoV-2 coronavirus – the causative agent of the COVID-19 (Coronavirus disease-2019) pandemic – are under intense current scrutiny, to understand how the virus operates and to uncover ways in which the disease can be prevented or alleviated.

Recent cell-based analyses of SARS-CoV-2 protein - protein interactions have mapped the human proteins targeted by the virus. The DNA polymerase α - primase complex or primosome – responsible for initiating DNA synthesis in genomic duplication – was identified as a target of nsp1 (non structural protein 1), a major virulence factor in the SARS-CoV-2 infection.

Here, we report the biochemical characterisation of the interaction between nsp1 and the primosome and the cryoEM structure of the primosome - nsp1 complex. Our data provide a structural basis for the reported interaction between the primosome and nsp1. They suggest that Pol α - primase plays a part in the immune response to the viral infection, and that its targeting by SARS-CoV-2 aims to interfere with such function.

## Introduction

The enormous disruption to public health and the global economy caused by COVID-19 has spurred international research efforts to unravel the molecular biology of the SARS-CoV-2 coronavirus, in order to devise effective means of therapeutic and prophylactic intervention.

Coronaviruses comprise a large family of positive-sense single-strand RNA viruses that cause respiratory and enteric disease in animals and humans (Artika et al., 2020; Hartenian et al., 2020; Wang et al., 2020). Seven coronaviruses that are capable of infecting humans have been identified to date, belonging to the genera *alphacoronavirus* (HCoV-229E, HCoV-NL63) and *betacoronavirus* (HCoV-OC43, HCoV-HKU1, MERS, SARS-CoV, SARS-CoV-2) (Cui et al., 2019; Fung & Liu, 2019). Within the *betacoronavirus* genus, HCoV-OC43 and HCoV-HKU1 generally cause only a mild illness; in contrast, infection with MERS-CoV, SARS-CoV and SARS-CoV-2 results in a severe respiratory disease in some patients, with significantly increased fatality rates.

SARS-CoV-2 is most closely related to SARS-like coronaviruses from horseshoe bats found within China; it shares ∼79% of its sequence with SARS-CoV and ∼50% with MERS-CoV. Since its initial identification and characterisation (Wu et al., 2020; Zhou et al., 2020), genomic sequencing has shown that the SARS-CoV-2 genome encodes – from its 5’ to the 3’-end – two large polyproteins 1a and 1ab that are cleaved by viral proteases to generate 16 non-structural proteins (nsp1 - nsp16), four structural proteins (S, E, M, N) and a number of other accessory proteins.

Upon viral infection, the host cell mounts a rapid response of its innate immune system via the interferon (IFN) pathway, resulting in the secretion of type-I and type-III interferons and subsequent activation of a signalling pathway that leads to the expression of a wide-range of IFN-stimulated genes (Lazear et al., 2019). A hallmark of COVID-19 is the attenuation of the interferon response of the host coupled to high levels of inflammatory cytokine production (Blanco-Melo et al., 2020). SARS-CoV-2 deploys multiple mechanisms to suppress the innate immune response, with participation of several viral proteins including nsp1, nsp6, nsp13, M, ORF3a, ORF6 and ORF7a/b (Lei et al., 2020; Xia et al., 2020).

The nsp1 protein is present in α- and β-coronaviruses, but not in δ- or γ-coronaviruses. nsp1 is considered to be a major virulence factor for SARS and MERS, due its pleiotropic activities as potent antagonist of virus- and IFN-dependent signalling and suppressor of host mRNA translation (Kamitani et al., 2006; Narayanan et al., 2008; Terada et al., 2017; Wathelet et al., 2007). While there is poor sequence conservation between the nsp1 proteins from α- and β-coronaviruses, both have been shown to induce strong suppression of host gene expression, albeit possibly by different mechanisms (Kamitani et al., 2009; Lokugamage et al., 2015; Narayanan et al., 2015; Shen et al., 2019). Despite poor sequence identity, α-CoV nsp1 and β-CoV nsp1 share structural homology in their core domain, which folds in a 6-stranded β-barrel structure with one α-helix on the rim of the barrel (Almeida et al., 2007; Clark et al., 2020; Jansson, 2013; Semper et al., 2021; Shen et al., 2018). The importance and role of nsp1’s globular domain is not yet fully understood, although it has recently been shown to interact with the 5′-UTR of the SARS-CoV-2 mRNA, helping it escape translational inhibition in infected cells (Shi et al., 2020). In addition to their globular domain, SARS-CoV and SARS-CoV-2 nsp1 proteins contain a long, unstructured C-terminal extension. A two-helix hairpin motif near the end of the C-terminal tail of SARS-CoV-2 nsp1 has been shown to bind the 40S subunit of the ribosome and block the mRNA entry channel, providing a structural basis for the role of nsp1 in suppressing host translation in infected cells (Kamitani et al., 2009; Lapointe et al., 2021; Schubert et al., 2020; Thoms et al., 2020).

Recently published studies of the human proteins targeted by SARS-CoV-2 reported a physical association between nsp1 and the primosome (Gordon, Hiatt, et al., 2020; Gordon, Jang, et al., 2020), the complex of DNA polymerase α (Pol α) and primase responsible for initiating DNA synthesis in DNA replication (Pellegrini, 2012). In addition to its well-characterised nuclear role in genomic duplication, participation of Pol α in innate immunity has been suggested, based on the observation that disease-associated mis-splicing of *POLA1* – coding for the catalytic subunit of Pol α – results in abnormal interferon I response and autoinflammatory manifestations (Esch et al., 2019; Starokadomskyy et al., 2016, 2021). Here, we demonstrate that nsp1 binds directly to the catalytic subunit of Pol α and elucidate their mode of interaction by determining the cryo-EM structure of SARS-CoV nsp1 bound to the human primosome. Our biochemical and structural characterisation of the interaction between the virulence factor nsp1 and the primosome suggests that targeting Pol α - primase is part of a novel mechanism driving the SARS-CoV-2 infection. Our data further define protein epitopes on nsp1 and Pol α catalytic subunit that could be exploited for the development of small-molecule inhibitors of their interaction.

## RESULTS

### Biochemical analysis of the nsp1-primosome interaction

A proteome-wide SARS-CoV-2 protein interaction study, performed by co-immunoprecipitation of exogenously expressed Strep-tagged SARS-CoV-2 proteins from HEK293T cells, identified the four components of the Pol α - primase complex as high-confidence interactors of SARS-CoV-2 nsp1 (Gordon, Jang, et al., 2020). A subsequent study, which expanded this approach to include the SARS-CoV and MERS protein-protein interaction networks, similarly identified the four primosome components as SARS-CoV nsp1 interactors (Gordon, Hiatt, et al., 2020). We therefore initially sought to confirm a direct interaction between purified, recombinant samples of nsp1 and the human primosome.

MBP-tagged SARS-CoV-2 and SARS-CoV nsp1 were able to pull-down both full-length and truncated forms of purified human primosome (Fig. 1A). The truncated primosome lacks the flexible and largely unstructured N-terminal regions of both Pol α (amino acids 1 - 333) and B subunit (amino acids 1 - 148), which are known to mediate several primosome interactions (Evrin et al., 2018; Kilkenny et al., 2017; Kolinjivadi et al., 2017; Simon et al., 2014), thus indicating that the interaction with nsp1 is not dependent on known recognition motifs in Pol α. MERS nsp1 was also able to interact with the primosome, albeit more weakly than the SARS nsp1 proteins (Fig. 1B), in accordance with the proteome-wide interaction studies that identified the primosome as a high-confidence interactor of SARS-CoV and SARS-CoV-2, but not of MERS nsp1 (Gordon, Hiatt, et al., 2020). Importantly, nsp1 from human-infective coronaviruses strains 229E, NL63 and HKU1 did not interact with the primosome in our pull-down assay (Fig 1C).

**Figure 1.**
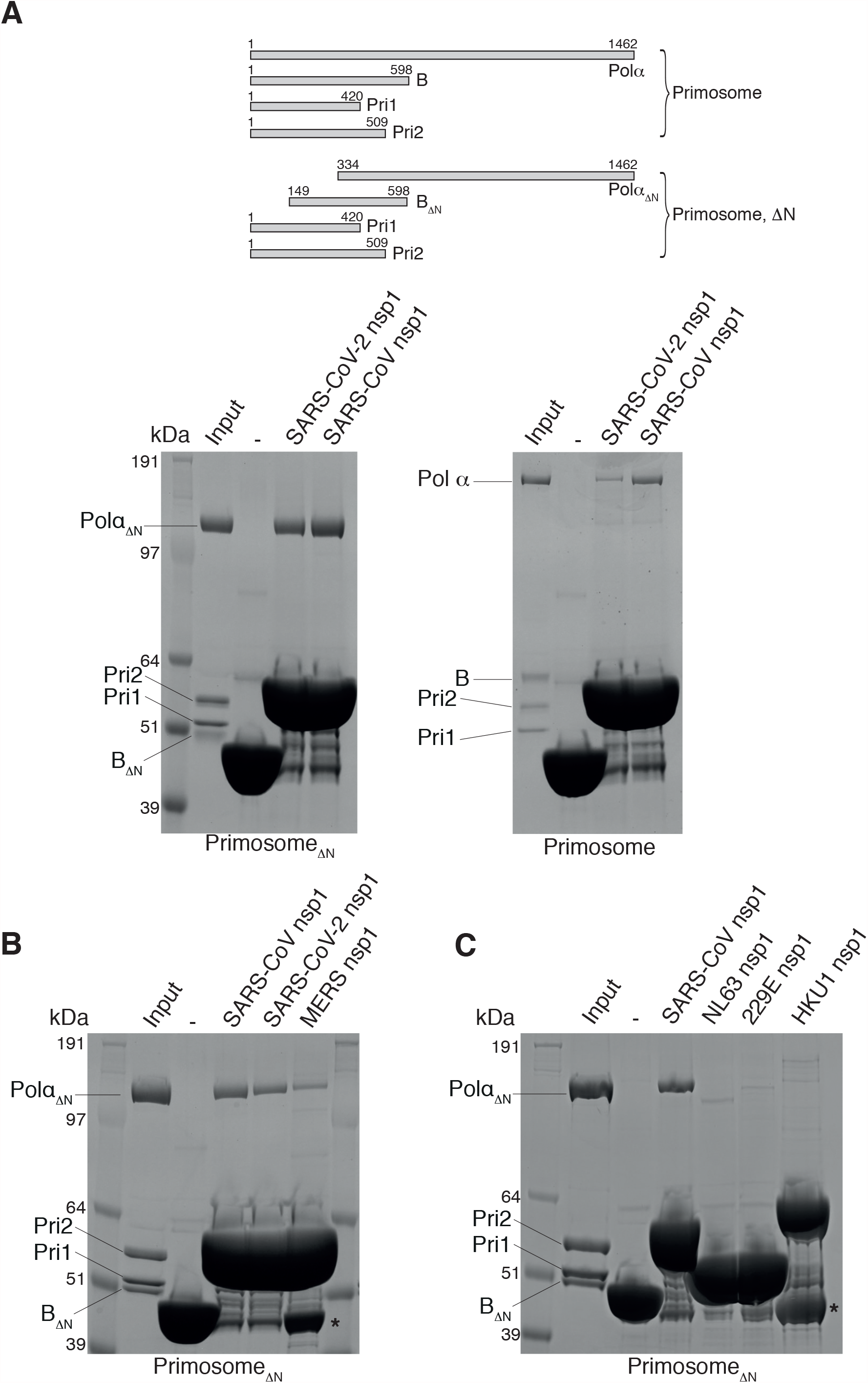
Biochemical reconstitution of the primosome - nsp1 interaction. The panels report the SDS-PAGE analyses of pull-down experiments on amylose beads of human primosome by maltose-binding protein (MBP) fusions of nsp1 proteins. **A** Pull-down of full-length and truncated primosomes. On the right, schematic drawing of subunit size for full-length and truncated primosomes. **B** Pull-down of truncated primosome with SARS-CoV, SARS-CoV-2 and MERS nsp1. **C** Pull-down of truncated primosome with SARS-CoV, NLS63, 229E and HKU1 nsp1. Bands labelled with an asterisk indicate free His_6_-MBP.

In the original interaction study (Gordon, Jang, et al., 2020), nsp1 was co-precipitated with all four subunits of Pol α - primase, which are normally found associated as a constitutive complex in the cell. To dissect the interaction between primosome components and SARS-CoV-2 nsp1, we performed a pull-down experiment using MBP-tagged nsp1 and the primase heterodimer Pri1 - Pri2 (Fig. 2A). We detected no interaction between nsp1 and primase which, taken together with the data presented in figure 1A, indicates that the nsp1-binding site resides within the folded domains of Pol α and the B subunit.

**Figure 2.**
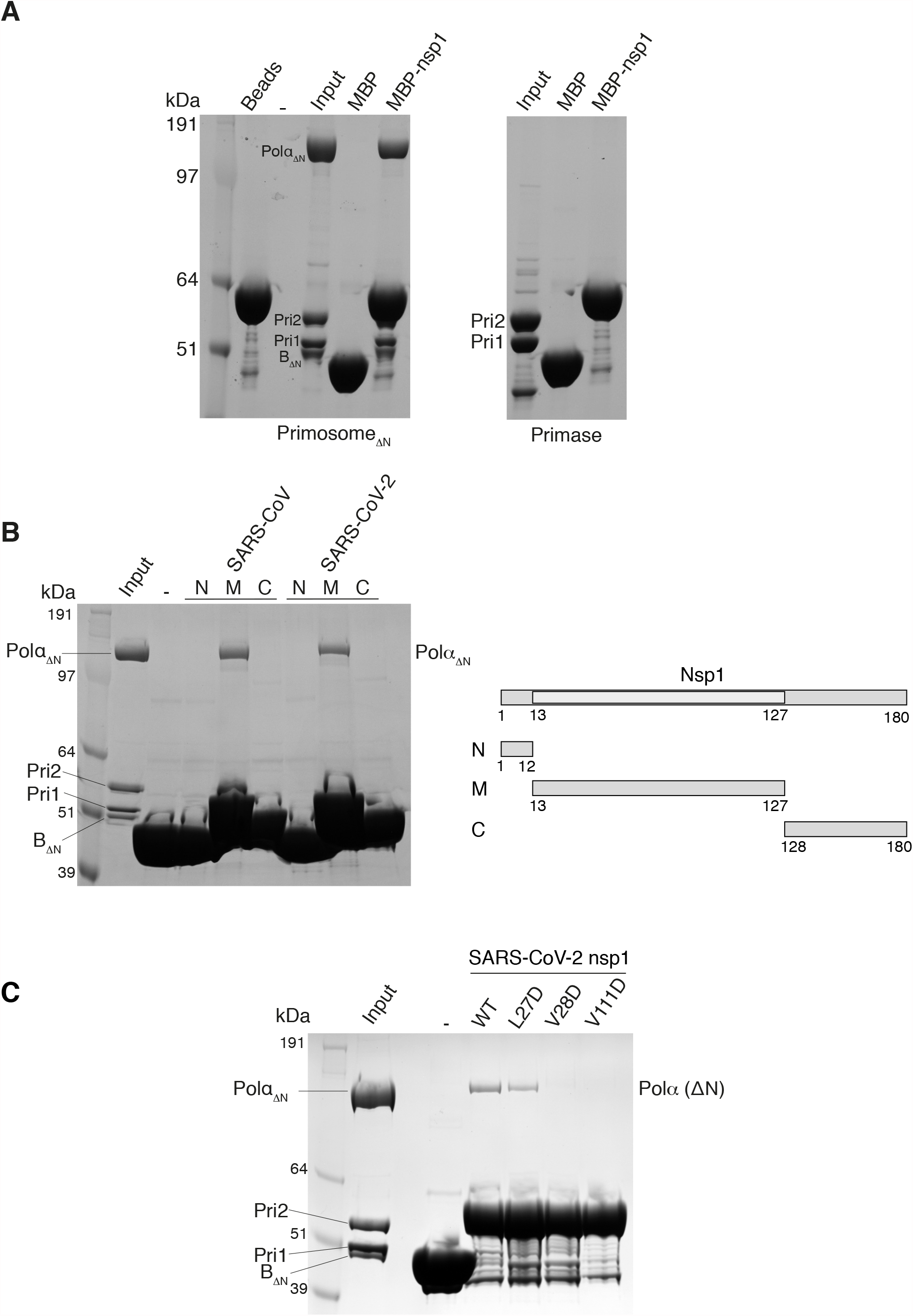
Biochemical dissection of the primosome - nsp1 interaction. The panels report the SDS-PAGE analysis of pull-down experiments by maltose-binding protein (MBP) fusions of nsp1 proteins immobilised on amylose beads. **A** Pull-down of truncated primosome and primase. **B** Pull-down of truncated primosome by N-, M- and C-terminal domains of SARS-CoV and SARS-CoV-2 nsp1. On the right, schematic diagram of the domain structure of nsp1. **C** Pull-down of truncated primosome by wild-type and single-point mutant SARS-CoV-2 nsp1.

Next, we sought to define the region of nsp1 responsible for the interaction with the primosome. SARS-CoV and SARS-CoV-2 nsp1 folds into a globular middle domain (M; amino acids 13 - 127) flanked by a short N-terminal region (N; amino acids 1-12), and a long C-terminal extension that is predicted to be largely unstructured (C; amino acids 128 - 180). Pull-down experiments with the N, M and C regions of SARS nsp1 showed clearly that the middle domain is responsible for the interaction with the primosome (Fig. 2B).

Guided by the published atomic model of SARS-CoV nsp1 (Almeida et al., 2007), we identified an exposed hydrophobic surface centred around amino acids L27 and V28 in the first inter-strand loop of the β-barrel in the middle domain and V111 in the last inter-strand loop, as a possible primosome-binding epitope. Point mutations to aspartate – that reversed the chemical nature of the side chain – weakened (L27D) or abolished (V28D and V111D) the interaction of the MBP-tagged nsp1 mutants with the primosome in our pull-down assay (Fig. 2C), thus implicating these residues in the interaction.

### CryoEM structure of the primosome - nsp1 complex

Our findings had shown that a direct biochemical interaction exists between nsp1 and the primosome, which was most likely the determinant of their co-precipitation in human cells (Gordon, Hiatt, et al., 2020; Gordon, Jang, et al., 2020). We therefore decided to elucidate the structural basis for the interaction by determining the high-resolution structure of the primosome - nsp1 complex by cryoEM (Supplementary figs. 1 - 3 and Supplementary table 1). SARS-CoV nsp1 was chosen for the structural analysis as this variant showed somewhat stronger binding to human primosome in our pull-down assays (Fig. 1A). We note that there is a high degree of sequence conservation between SARS-CoV and SARS-CoV-2 nsp1 (84.4% and 85.8% identity over the entire sequence and the middle domain only, respectively).

We obtained a 3D reconstruction of the primosome at a global resolution of 3.8 Å after sharpening (0.143 FSC) that showed clearly all components of the complex (Supplementary figs. 4 - 5). As the portion of the map relative to the primase subunit Pri1 was weaker, a new map was calculated using subtracted particles from which Pri1 had been omitted, yielding a slightly improved map at a global resolution of 3.6 Å after sharpening (0.143 FSC). Both maps – from full and subtracted particles – were used during model building. Density belonging to the nsp1 middle domain was clearly visible, at a lower map threshold than required to visualise the primosome core.

#### Primosome conformation

The cryoEM structure of the human primosome confirmed the presence of the binary interfaces that had been previously defined by X-ray crystallography of yeast and human Pol α - primase subcomplexes (Baranovskiy et al., 2015; Kilkenny et al., 2012, 2013; Klinge et al., 2009; Suwa et al., 2015). Intriguingly, our cryoEM analysis of the primosome revealed a quaternary structure that is incompatible with known and presumed conformations required for activity by its two catalytic subunits, Pol α and primase (Baranovskiy et al., 2014; Coloma et al., 2016; Kilkenny et al., 2013; Perera et al., 2013) (Fig. 3). Thus, the B subunit bound to the C-terminal domain of Pol α is docked within the open ‘hand’ of the polymerase domain, precluding binding of the template DNA - primer RNA duplex. Furthermore, the heterodimeric primase is held is an extended conformation, whereby the Fe-S domain of Pri2 – required for initiation of RNA primer synthesis – is sequestered by interactions with both B and Pol α and unavailable to contact Pri1, which is excluded from the primosome core and largely solvent exposed.

**Figure 3.**
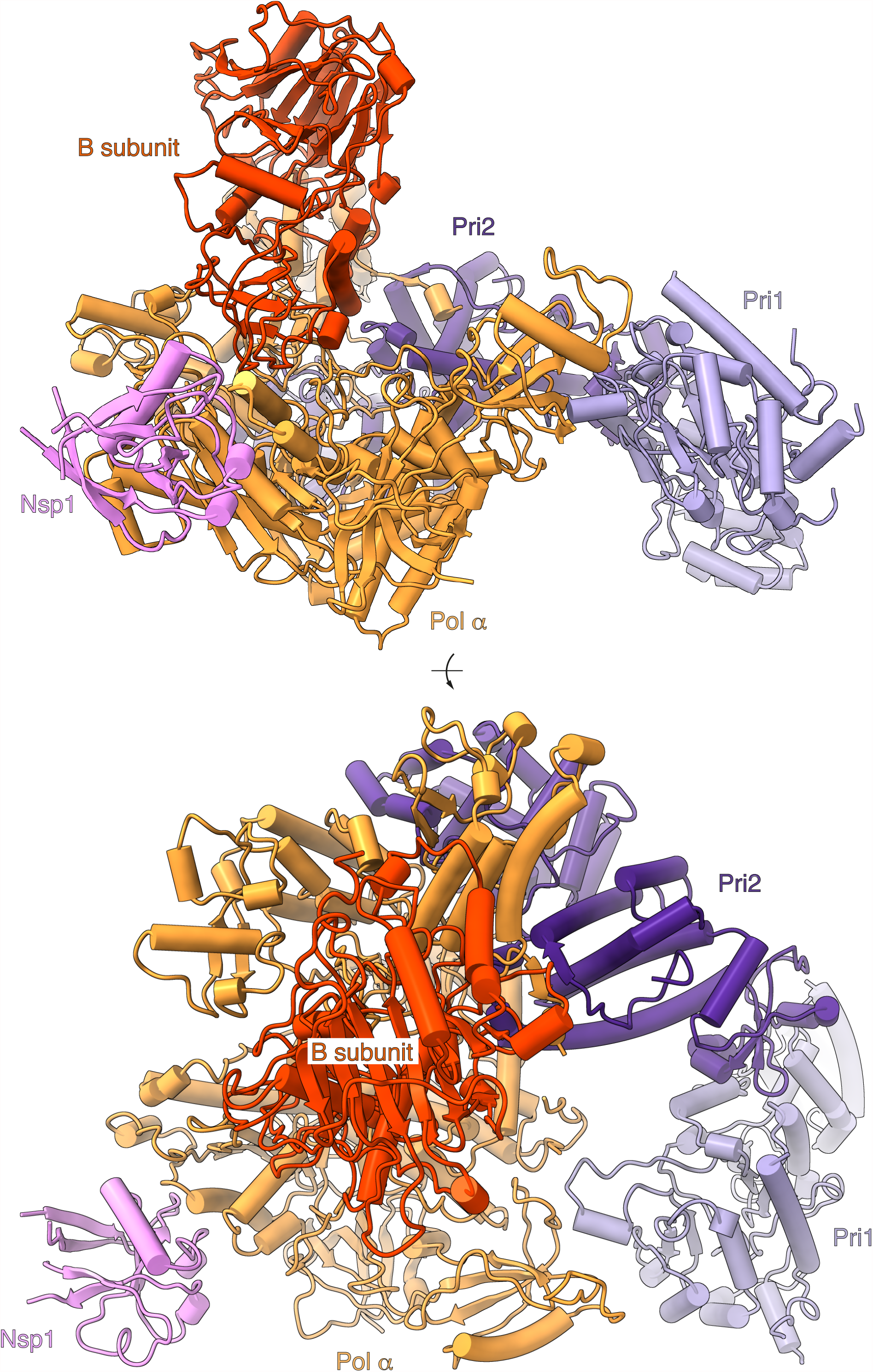
CryoEM structure of the primosome - nsp1 complex. The structure is drawn as a cartoon with helices as solid cylinders and strands as arrows. Each subunit is labelled and uniquely coloured, in orange (Pol α), red (B subunit), Pri1 (purple), lilac (Pri2) and pink (nsp1). The structure is shown in two views, from the side (top panel) and from the top (bottom panel).

Overall, the cryoEM structure is highly similar to a structure of the human primosome determined earlier by X-ray crystallography (Baranovskiy et al., 2016) (Supplementary fig. 6). Although reasonable doubts existed that the crystal structure might have been a crystallisation artefact, these have now been dispelled by our cryoEM analysis. Altogether, the high-resolution structure data obtained with X-ray and cryoEM methods indicate that – in its unliganded form – the human primosome exists in a conformation that is incompatible with primer synthesis without the intervention of major conformational rearrangements. Why should the primosome adopt such a distinct quaternary structure in the absence of its DNA and nucleotide substrates and what should be the function of this postulated autoinhibited state (Baranovskiy et al., 2016) remains unclear.

#### Interaction with nsp1

The structure shows that nsp1 binds to the Pol α subunit of the primosome (Fig. 4). The globular domain of nsp1 docks onto the rim of the polymerase fold of Pol α, and its binding site is entirely contained within the exonuclease domain of the polymerase, in agreement with our biochemical findings (Figs. 1 and 2). Nsp1 binding is mediated by residues in the first, second and fourth inter-strand loop of the globular β-barrel domain, and in the α-helix at the C-end of the second inter-strand loop. The interface is of a mixed hydrophobic and polar nature (Fig. 5). A cluster of hydrophobic nsp1 residues at the centre of the interface – including L27, V28, P109, V111, G112 – becomes buried upon complex formation (Fig. 5A and Supplementary fig. 7), validating the observation that mutations of nsp1 residues V28 and V111 abrogate the interaction (Fig. 2C).

**Figure 4.**
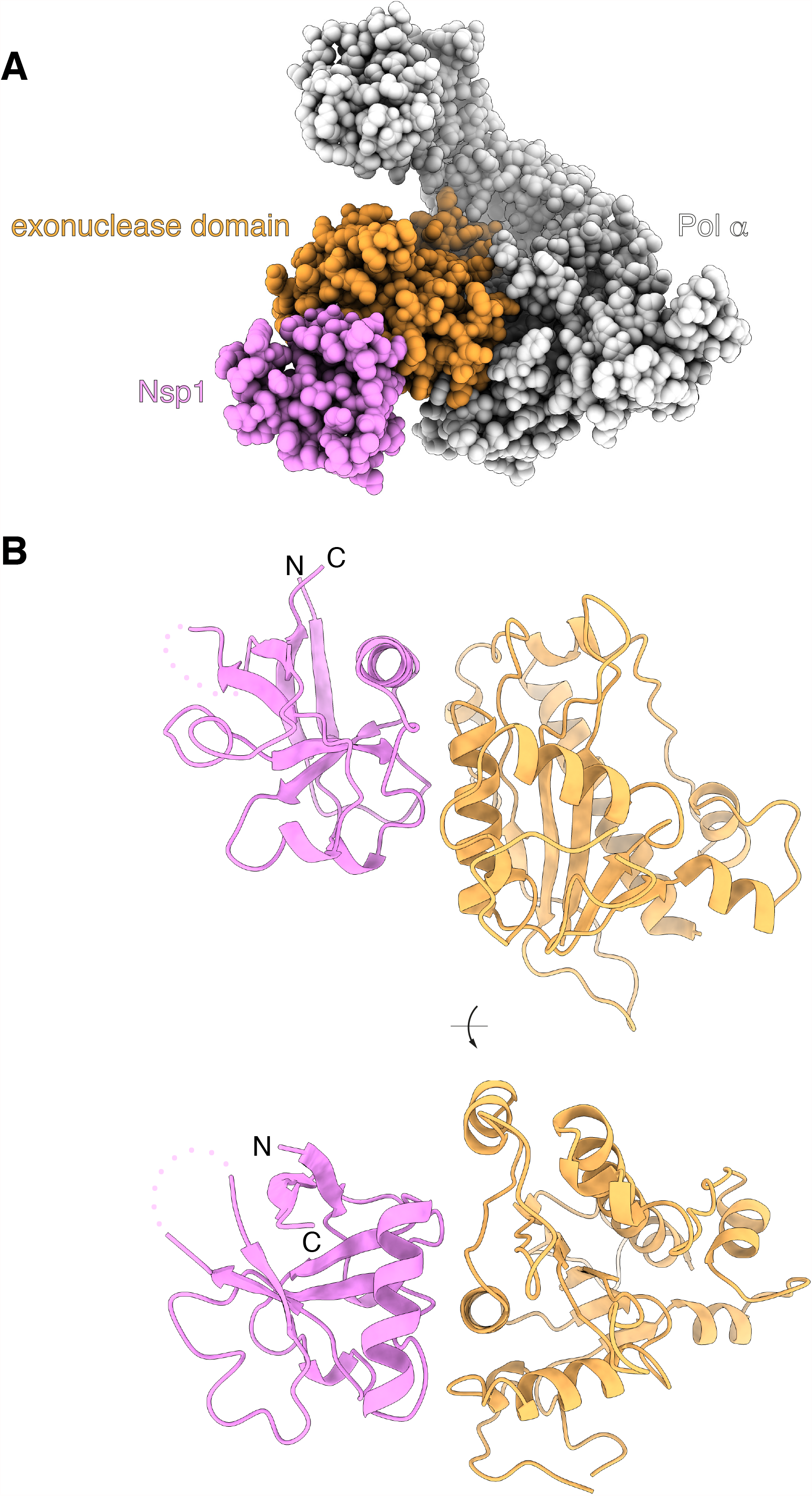
nsp1 binds to the exonuclease domain of Pol α. **A** Space-fill representation of the interaction between nsp1 (pink) and the inactive exonuclease domain of Pol α (orange). **B** Two views rotated by 90 degrees of the association mode of nsp1 with the exonuclease domain. The proteins are drawn as ribbons, in pink (nsp1) and orange (Pol α). A disordered loop of nsp1 facing away from the interface is shown as a dotted line.

**Figure 5.**
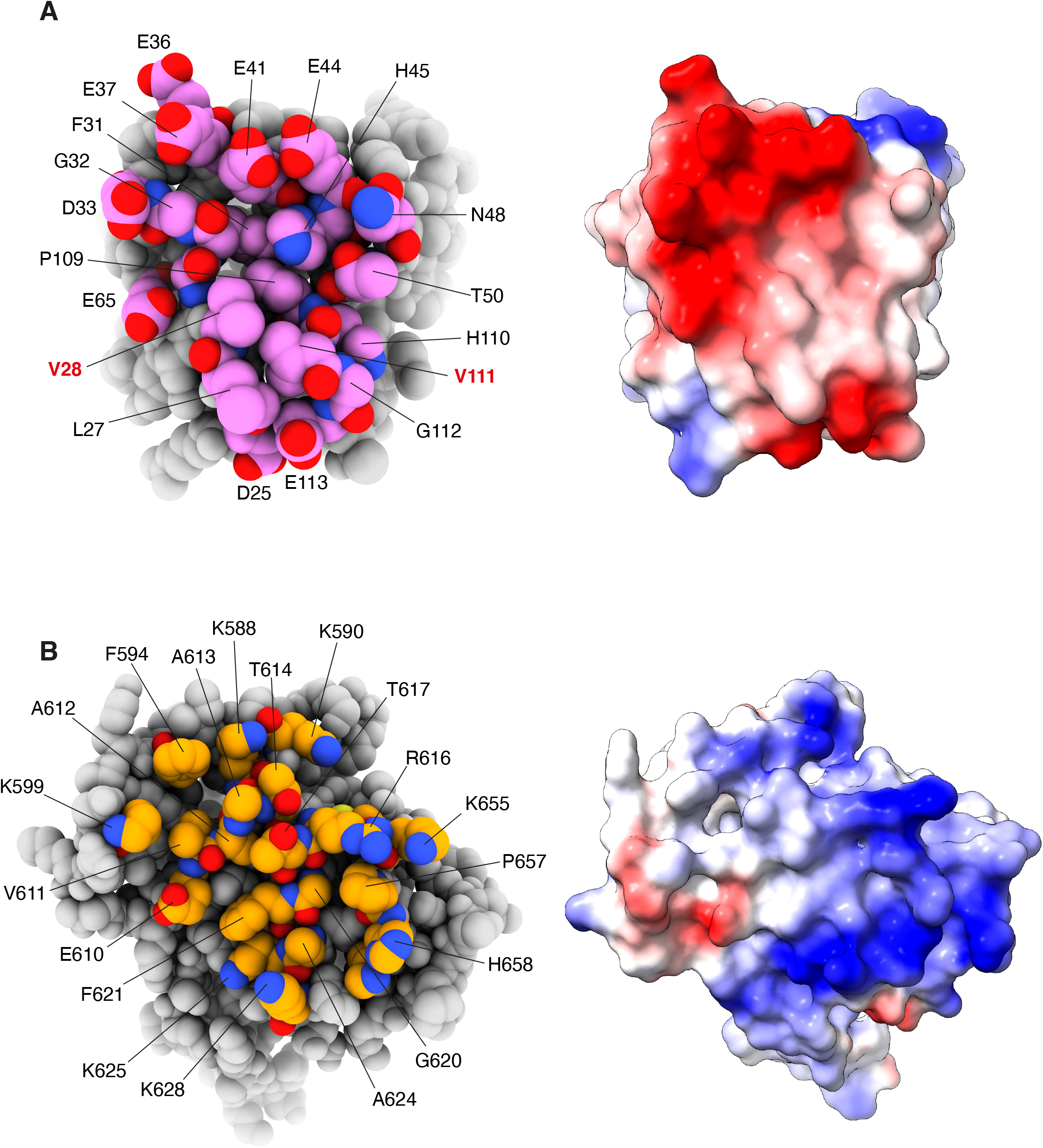
The nsp1 - Pol α exonuclease domain interface. **A** On the left, a space-fill model of nsp1, with interface residues labelled and coloured in pink. On the right, the charge distribution of nsp1 mapped over its molecular surface (red: negative charge; blue: positive charge). **B** On the left, a space-fill model of the exonuclease domain of Pol α, with interface residues labelled and coloured in orange. On the right, the charge distribution of the exonuclease domain mapped over its molecular surface (red: negative charge; blue: positive charge).

The nsp1-binding area in Pol α is centred on the second α-helix of the exonuclease domain, comprising residues 615 to 629 (recognition helix), and extends to include most of the sequence of the solvent-exposed half of the exonuclease domain, from V585 to I662 (Fig. 5B and Supplementary fig. 7). Hydrophobic amino acids in the exonuclease domain that match the hydrophobic surface of nsp1 in the complex are helical residues A624, G620 and F621, the short aliphatic segment 611-VAAT-614 in the loop leading into the recognition helix, and P657 towards the C-end of the binding interface.

The interaction shows pronounced charge complementarity (Fig. 5A, B and Supplementary fig. 7). A cluster of negatively charged residue in nsp1 – D33, E36, E37, E41 and E44 in the helix of the second intra-strand loop, as well as D25, E65 and E113 – lines the edge of the interaction area, and becomes juxtaposed upon complexation to basic residues K625, K628 and R616 in the recognition helix of the exonuclease domain of Pol α, as well as surrounding lysine residues K588, K590, K599, K655 and K661.

The presence at the interface of small amino acids such as alanine and glycine, that are required for the intimate association of rather flat protein surfaces, supports the observed interaction between nsp1 and Pol α. Notably, the nsp1-binding sequence 611-VAAT-614, as well as the 620-GFxxA-624 motif in the recognition helix, are highly conserved in vertebrate Pol α sequences (Supplementary fig. 7).

## Discussion

Here we have provided a biochemical and structural basis for the interaction between the SARS-CoV-2 virulence factor nsp1 and the primosome, for which evidence first emerged from protein - protein interaction screens between SARS-CoV-2 and human proteins. Our structure shows that nsp1 uses its globular domain to interact with the primosome, distinct from the C-terminal motif employed to target the ribosome and suppress mRNA translation. Thus, our findings provide the first structural evidence of how nsp1 can deploy its conserved middle domain to engage with a host protein.

The ability of nsp1 to use different regions of its sequence to target different physiological processes in the cell is in keeping with evidence of its pleiotropic pathological effects during infection. Furthermore, as viral proteins are known to exploit existing binding epitopes on the surface of the target protein, the nsp1-binding site in Pol α might be indicative of as-yet unknown physiological partners of the primosome binding at this site. The discovery of a novel protein-protein interaction surface in nsp1 and Pol α opens up the possibility of developing small-molecule inhibitors aimed at disrupting their association, which might of therapeutical value in the treatment of COVID-19.

How the interaction of nsp1 with the primosome might promote virulence is currently unclear. However, a rationale for targeting the primosome by SARS-CoV-2 might be envisaged, based on recent evidence that mutations in *POLA1* – coding for the catalytic subunit of Pol α – cause XLPDR (X-linked reticulate pigmentary disorder), a genetic syndrome that is characterised by activation of type I interferon signalling, a persistent inflammatory state and recurrent lung infections (Starokadomskyy et al., 2016, 2021). A similar set of symptoms is observed in individuals with inherited haploinsufficiency of the *PRIM1* gene, coding for the Pri1 subunit of primase (Parry et al., 2020). In XLPDR fibroblasts, the ability of Pol α - primase to synthesise RNA/DNA hybrids in the cytoplasm was found to be impaired (Starokadomskyy et al., 2016). This effect was proposed to be responsible for the disease, although the molecular mechanisms by which cytosolic RNA/DNA hybrids might help regulate the anti-viral response are unknown.

To test whether nsp1 binding may directly affect the enzymatic activity of the primosome, we performed biochemical assays to measure the RNA and DNA primer synthesis activities of primase and Pol α, respectively. However, we could not detect an effect on RNA primer synthesis or DNA extension in the presence of an excess of nsp1 protein (Supplementary fig. 8). This is consistent with the 3-D structure of the primosome-nsp1 complex, in which nsp1 is docked in a position that would not be predicted to interfere with RNA-DNA primer synthesis.

Given the exclusively cytoplasmic localisation of SARS and MERS nsp1 upon infection (Gordon, Hiatt, et al., 2020), the interaction with nsp1 appears aimed at targeting the primosome fraction present in the cytoplasm of infected cells. As nsp1 binding did not seem to impact the enzymatic activity of the primosome, we speculate that any effect on primosome activity might be achieved indirectly. Thus, nsp1 binding might be necessary for recruitment of additional factors that inhibit primosome activity or sequester it in a way that makes it inactive; alternatively, nsp1 binding might target the primosome for ubiquitination and degradation (Figure 6). More work will be required to establish the molecular mechanisms by which RNA/DNA synthesis by the primosome modulates the antiviral response of the host cell and how nsp1 binding affects this response.

**Figure 6.**
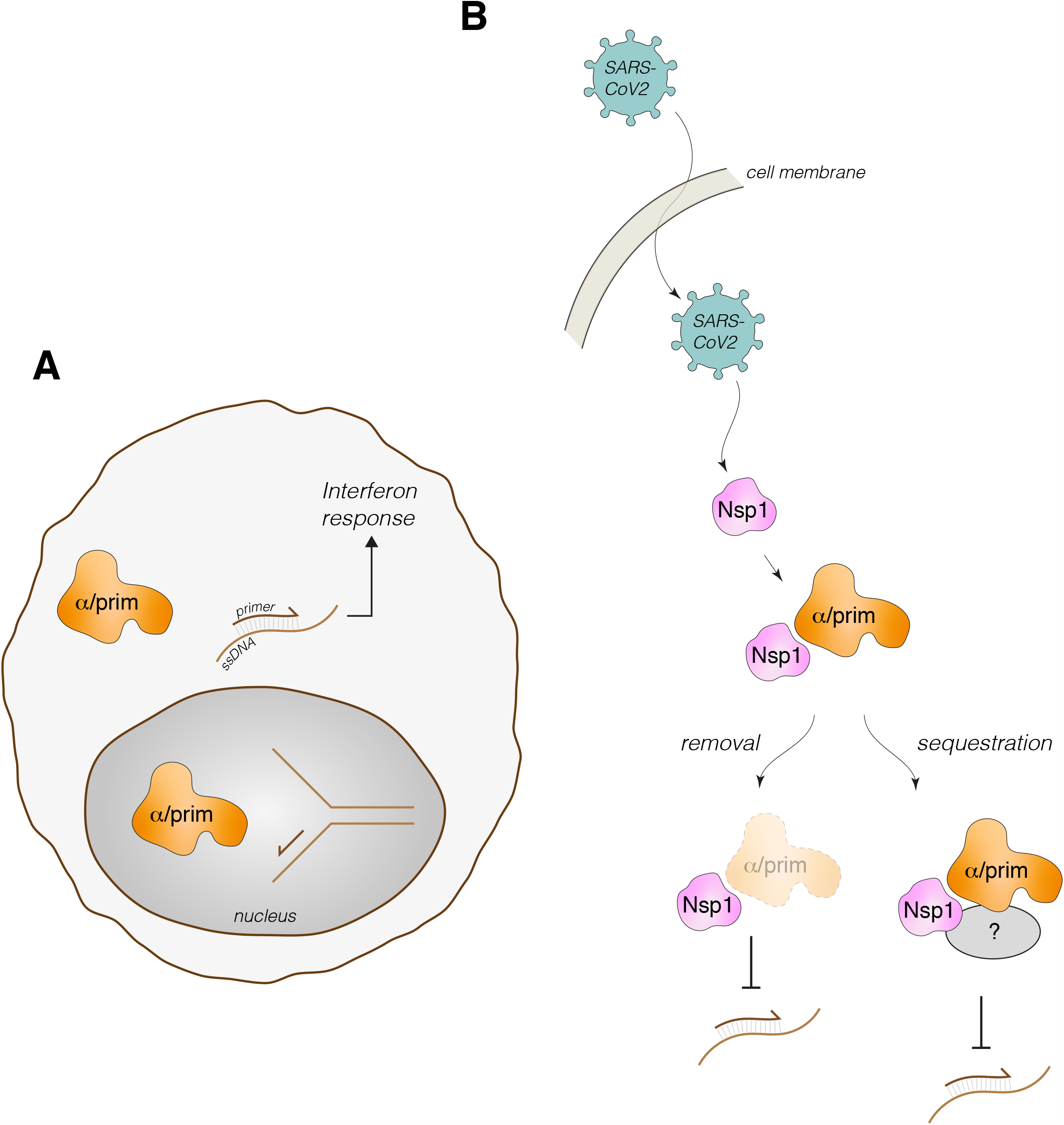
Possible functional significance of primosome targetting by SARS CoV2 nsp1. **A** In addition to its well-established role in chromosomal DNA duplication during S-phase, a cytoplasmic role for Pol α - primase has been postulated to synthesise RNA/DNA duplexes that might mediate the interferon response (Starokadomskyy et al., 2016). **B** Speculative mechanisms of nsp1-dependent SARS-CoV-2 interference with cytoplasmic Pol α - primase. Nsp1 binding might mediate sequestration of the primosome in an inactive form or promote its degradation.

## MATERIALS AND METHODS

### Cloning, expression and protein purification

Full-length human primosome (Pol α: 1-1462; B subunit: 1-598; PriS: 1-420; PriL: 1-509) and truncated human primosome (Pol α: 334-1462; B subunit: 149-598; PriS: 1-420; PriL: 1-509) was expressed in *Sf9* insect cells and purified as described previously (Kilkenny et al. 2017). Full-length coronavirus nsp1 genes (SARS-CoV-2, SARS-CoV, MERS, HKU1, NL63 and 229E) were ordered as synthetic gBlocks (IDT), and cloned into bacterial expression vector pMAT11 (Peranen et al., 1996). nsp1 proteins were expressed as His_6_-MBP-tagged fusion proteins from the Rosetta2 (DE3) *E. coli* strain (Novagen). nsp1 fragments (SARS-CoV N-terminus (1-12), middle domain (13-127), C-terminus (128-180); SARS-CoV-2 N-terminus (1-12), middle domain (13-127), C-terminus (128-180)) were cloned and expressed in the same way. Full-length SARS-CoV-2 nsp1 point mutants (V111D, L27D, V28D) were ordered as synthetic gBlocks (IDT), and cloned and expressed as above.

All His_6_-MBP-fusion proteins were purified by metal affinity chromatography, using Ni-NTA agarose (Qiagen). A high salt wash (500mM KCl) was performed on-column to remove any traces of nucleic acid from the protein preparation. Proteins were eluted using 25mM HEPES pH 7.2, 300 mM KCl and 200 mM imidazole.

### Cryo-EM sample preparation and grid freezing

SARS-CoV His_6_-MBP-nsp1 (13-127) was TEV-cleaved overnight at 4°C, and the His_6_-MBP tag was subsequently recaptured using amylose resin (NEB). The flow-through was loaded onto a Q-HP anion exchange column (Cytiva) and eluted with a KCl gradient (25 mM HEPES pH 7.2, 80-750 mM KCl, 2 mM DTT). Nsp1 and the truncated primosome were subsequently individually buffer-exchanged into cryo-EM buffer (25 mM HEPES pH 7.2, 150 mM KCl, 1 mM DTT) using a NAP5 column (Cytiva). The two proteins were mixed, at final concentrations of 1 μM primosome and 10 μM nsp1. 1mM BS3 crosslinker was added, and the sample incubated on ice while grid freezing commenced using a Vitrobot set at blot force −10 (Thermo Fisher Scientific). The samples were diluted 1:2 with cryo-EM buffer immediately before application of the protein sample to the grid. 3 μl sample was applied to each side of the UltrAuFoil grid (300 mesh, R 1.2/1.3, Quantifoil), which had been glow discharged on both sides (25 mA, 1 minute each side). The grids used for the eventual structure determination were plunge frozen in liquid ethane approximately 45 minutes after addition of the BS3 cross-linker.

### Cryo-EM data processing and structure refinement

Data collection parameters, structure refinement details and statistics are reported in Supplementary table 1 and Supplementary figure 1. 393 movies (1.11 e/Å^2^/fraction, 40 fractions) were collected on a Talos Arctica at 200keV, 92,000xmagnification and 1.13 px/Å, with a Falcon III detector in counting mode. Movie processing was carried out in Relion 3.1 (Scheres, 2012). The movies were motion corrected using Relion’s implementation and CTFs were estimated using CTFFIND4-1 (Rohou & Grigorieff, 2015). An initial set of 165,011 particles was auto-picked and used to generate 2D templates for template-based picking of a new set of 183,301 particles. The template-picked particle set was reduced to 97,680 particles by rounds of 2D classification and used to reconstruct an initial 3D volume. 3D classification into 4 models generated one model with 47,542 particles (48.6%) which was refined to 7.29 Å. The 3D volume showed clearly evidence of the four primosome subunits and of the bound nsp1 protein; the initial 3D model was used as starting point for 3D classification during processing of the higher-resolution dataset (see below).

2919 movies (0.98 e/Å^2^/fraction, 48 fractions) were collected on Titan Krios at 300 keV, 130,000 magnification and 0.326 Å px/Å with a K3 detector in super-resolution counting mode. Movie processing was carried out in Relion 3.1 (Scheres, 2012). The movies were motion corrected and 2xbinned using Relion’s own implementation and CTFs were estimated using CTFFIND4-1 (Rohou & Grigorieff, 2015). An initial set of 314,987 particles was auto-picked and improved by rounds of 2D classification, to generate 2D templates for template-based picking of a new set of 709,068 particles. The particles set was improved by multiple rounds of selection by 2D classification, and a first 3D classification into 4 models was performed using the initial 3D model obtained for the Talos dataset. Class #1 (251,901 particles) was selected and a second round of 3D classification was performed, leading to a new 3D class #4 with 233,476 particles. 3D refinement of this class yielded a map at 4.12 Å, which was sharpened in Phenix (Adams et al., 2010) using ‘phenix.local_aniso_sharpen’ to a resolution of 3.8 Å (0.143 FSC) and used for model interpretation and model building.

For model building, the crystal structure of human Pol α - primase (PDB ID 5EXR) was fitted manually in the map with ChimeraX (Goddard et al., 2018) and real-space refined in Phenix (Adams et al., 2010), together with local manual rebuilding in Coot (Emsley & Cowtan, 2004). Additional density corresponding to the known structure of nsp1 was clearly detectable, although at a lower map threshold, bound to Pol α. A model of SARS nsp1 was built based on PDB entries 7K3N, 2HSX and docked into the map in ChimeraX (Goddard et al., 2018). The Pol α primase - nsp1 complex was further real-space refined in Phenix (Adams et al., 2010).

Inspection of the 2D/3D classes and of the 3D refined map for the Pol α - primase - nsp1 complex showed that the density corresponding to the Pri1 subunit of primase was weaker. This observation was explained by the looser contact of Pri1 with the rest of the Pol α - primase, which was primarily mediated by the known interface with Pri2. To improve the map of the rest of Pol α - primase and of the bound nsp1, a mask was prepared in ChimeraX (Goddard et al., 2018) that excluded the Pri1 lobe and used to generate a set of subtracted particles from the 233,476 particles set used to generate the original 3D refined map. A new initial 3D model was first generated, and after a round of 3D classification class #3 (103,763 particles) was selected. 3D refinement of this class generated a 4.40 Å map, that was sharpened using ‘phenix.local_aniso_sharpen’ to a resolution of 3.6 Å (01.43 FSC) and used for model interpretation and model building. The map showed slightly better high-resolution features for the primosome core and clearer features for the nsp1 fold, and was therefore used as a visual aid in the refinement of the primosome - nsp1 complex structure.

### Pull-downs

150 μl amylose resin (NEB) was equilibrated in pull-down buffer (1x PBS, 5% glycerol, 0.1% Igepal CA-630, 1 mM TCEP or DTT). An excess of His_6_-MBP-nsp1 was added to saturate the beads with bait protein. The His_6_-MBP tag protein was used as negative control. Beads were washed twice with pull-down buffer to remove excess, unbound bait protein. BSA was added to a final concentration of 2.5 mg.ml^-1^. Primosome (full-length or truncated) was added to the beads, which were subsequently incubated at 4 °C for 1 hour with rolling. The beads were washed four times with 1ml pull-down buffer. Bound protein was eluted using 150 μl pull-down buffer supplemented with 50 mM maltose. Samples were analysed by SDS-PAGE.

### Primosome activity assay

Full-length His_6_-MBP-nsp1 (from SARS-CoV-2 and SARS-CoV) was TEV-cleaved overnight at 4°C. The His_6_-MBP tag was recaptured using amylose resin (NEB). The flow-through was loaded onto a Q-HP anion exchange column (Cytiva) and nsp1 was eluted with a KCl gradient (25 mM HEPES pH 7.2, 80-750 mM KCl, 2 mM DTT). Pure nsp1 was subsequently buffer exchanged into 25 mM HEPES pH 7.2, 100 mM KCl, 1 mM DTT using a NAP5 column (Cytiva).

An activity assay, to monitor the ability of Pol α to extend an RNA primer, was performed using a pre-annealed DNA-RNA template (DNA45: 5’-TTCTTT ATCTCTTATTTCCTTCTATTTTCCACGCGCCTTCTATTT-3’; RNA13: 5’-[6FAM]-GGCGCGUGGAAAA-3’, Sigma Aldrich). The reaction buffer comprised 25 mM HEPES pH 7.2, 100 mM KCl, 10 mM Mg(OAc)_2_ and 1 mM DTT. Each 100 μl reaction contained 2 μM DNA-RNA template, 1 mM dNTP, 10 μM nsp1 and 0.1 μM full-length primosome. Reactions were incubated at 25 °C, and 30 μl aliquots removed and quenched (by adding an equal volume of 95% formamide and 25 mM EDTA) at the indicated time points. Samples were heated to 70°C for 2 min, then loaded onto a 19% urea-polyacrylamide gel (running conditions: 450 V, 90 min, 0.5× TBE buffer). Gels were initially imaged by laser scanning (473 nm laser, Typhoon FLA 9000, GE Healthcare), to detect only the 6FAM-labelled products. Gels were subsequently stained using a 1:10000 dilution of SYBR Gold Stain (Thermo Fisher Scientific) in 0.5× TBE buffer, and re-imaged in the same way to detect all nucleic acid.

## Acknowledgements

We would like to thank Ed Greenwood, Thomas Crozier and Paul Lehner (Cambridge Institute of Therapeutic Immunology and Infectious Disease) for helpful discussions. This work was funded by a Wellcome Trust Investigator award (104641/Z/14/Z) to LP.

## Data availability

Coordinates and 3D reconstructions for the primosome - nsp1 complex have been deposited under accession codes 7OPL (PDB), 13020 and 13021 (EMDB).

