## Supplementary information for "Structural basis for the interaction of SARS-CoV-2 virulence factor nsp1 with Pol α - Primase"

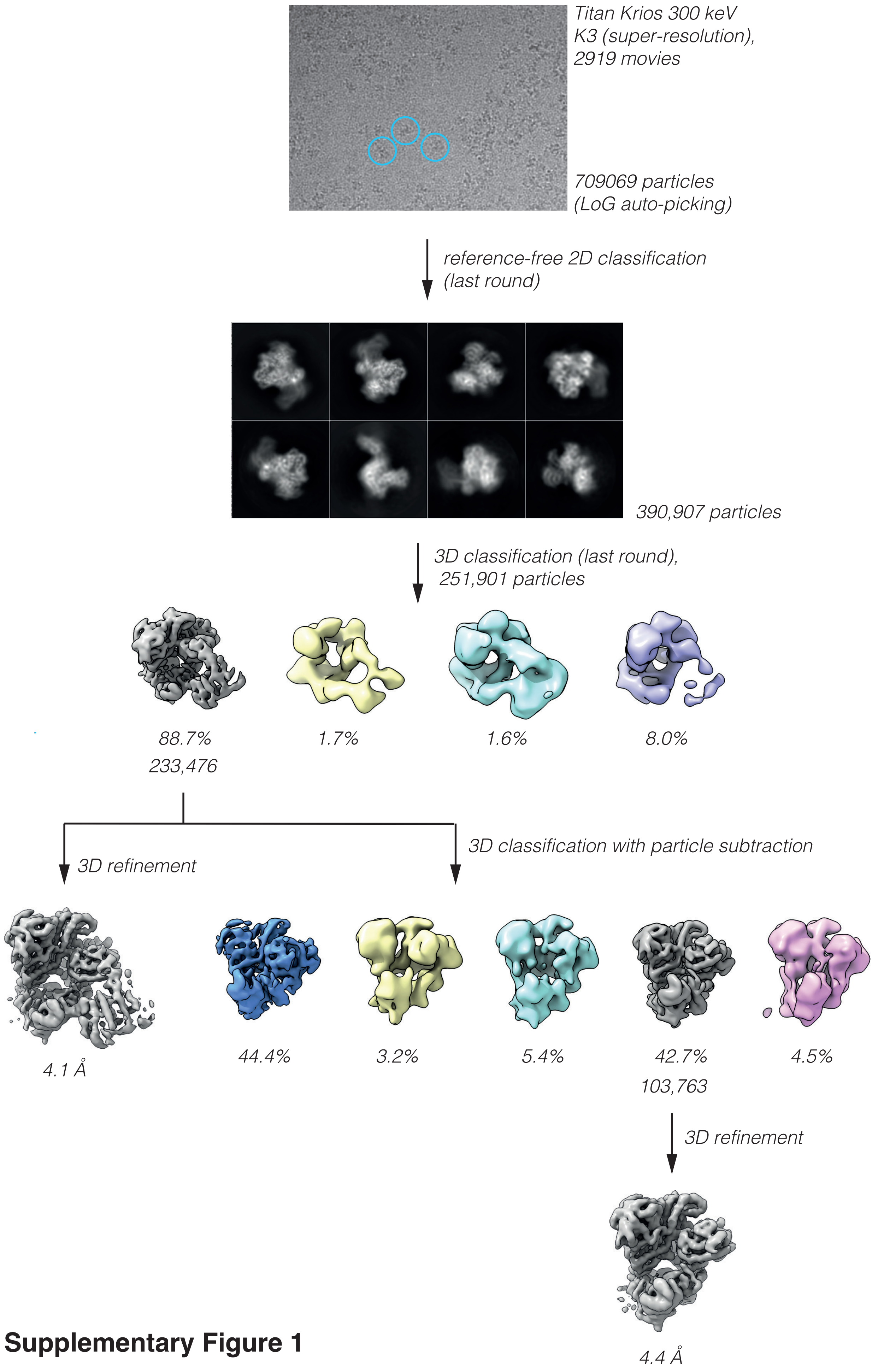

**Supplementary Figure 1**

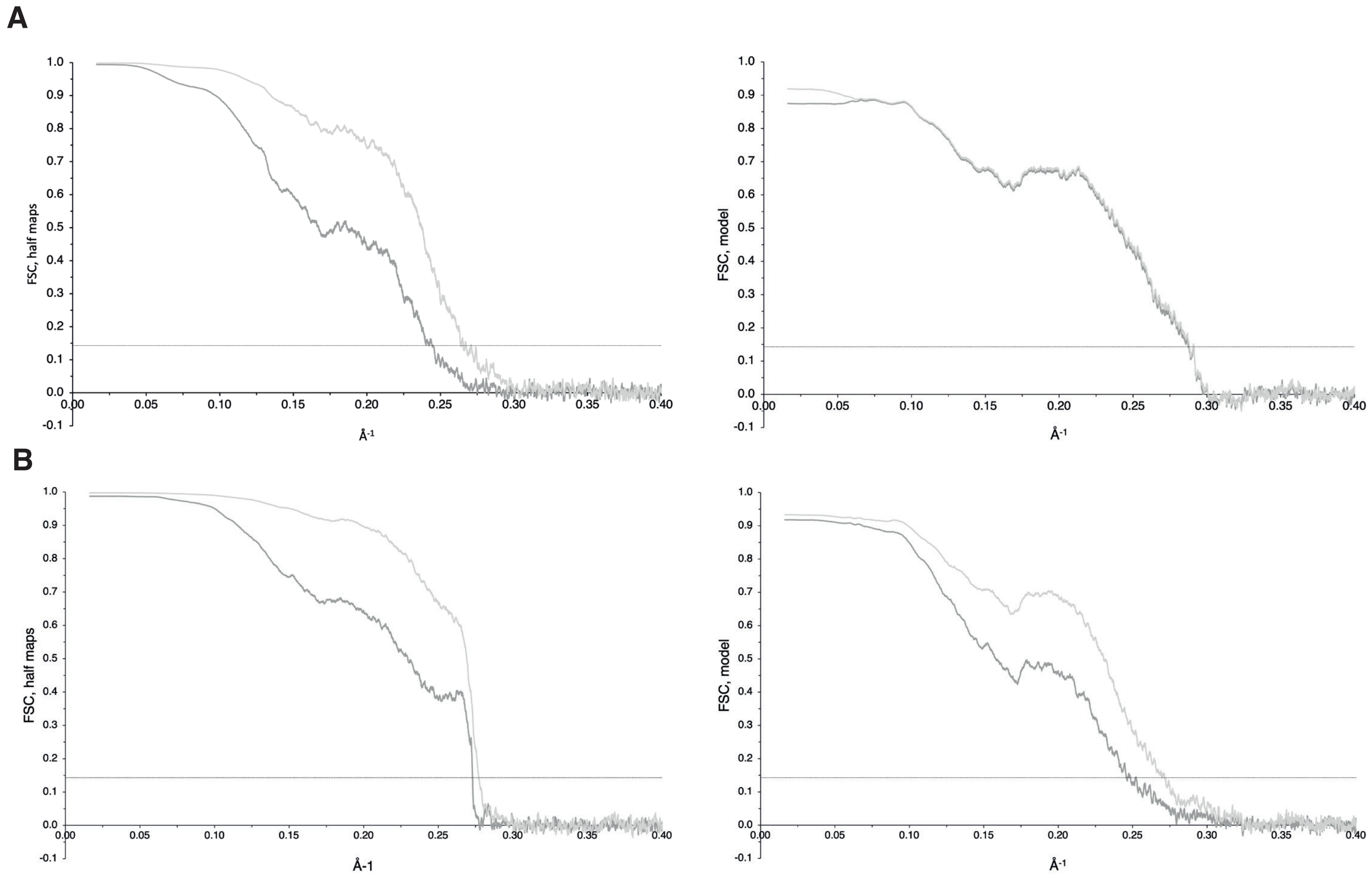

**Supplementary Figure 2**

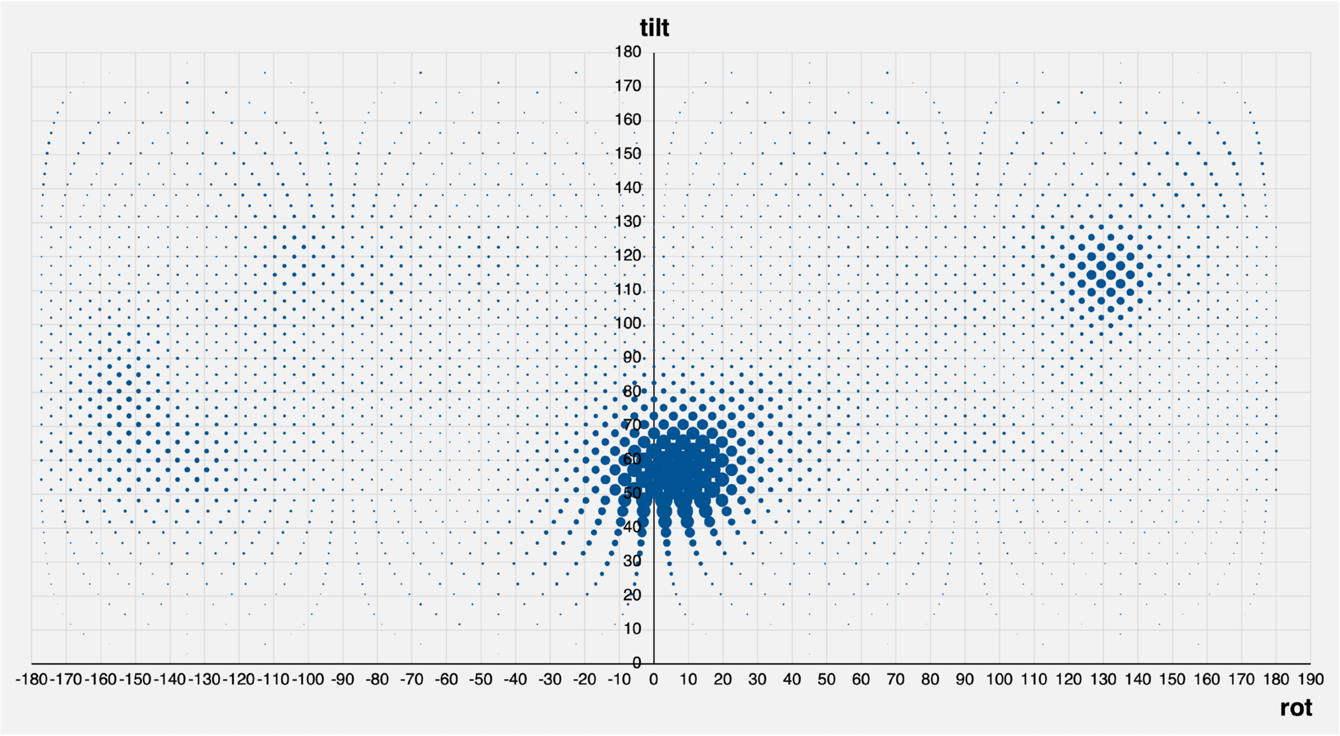

**Supplementary Figure 3**

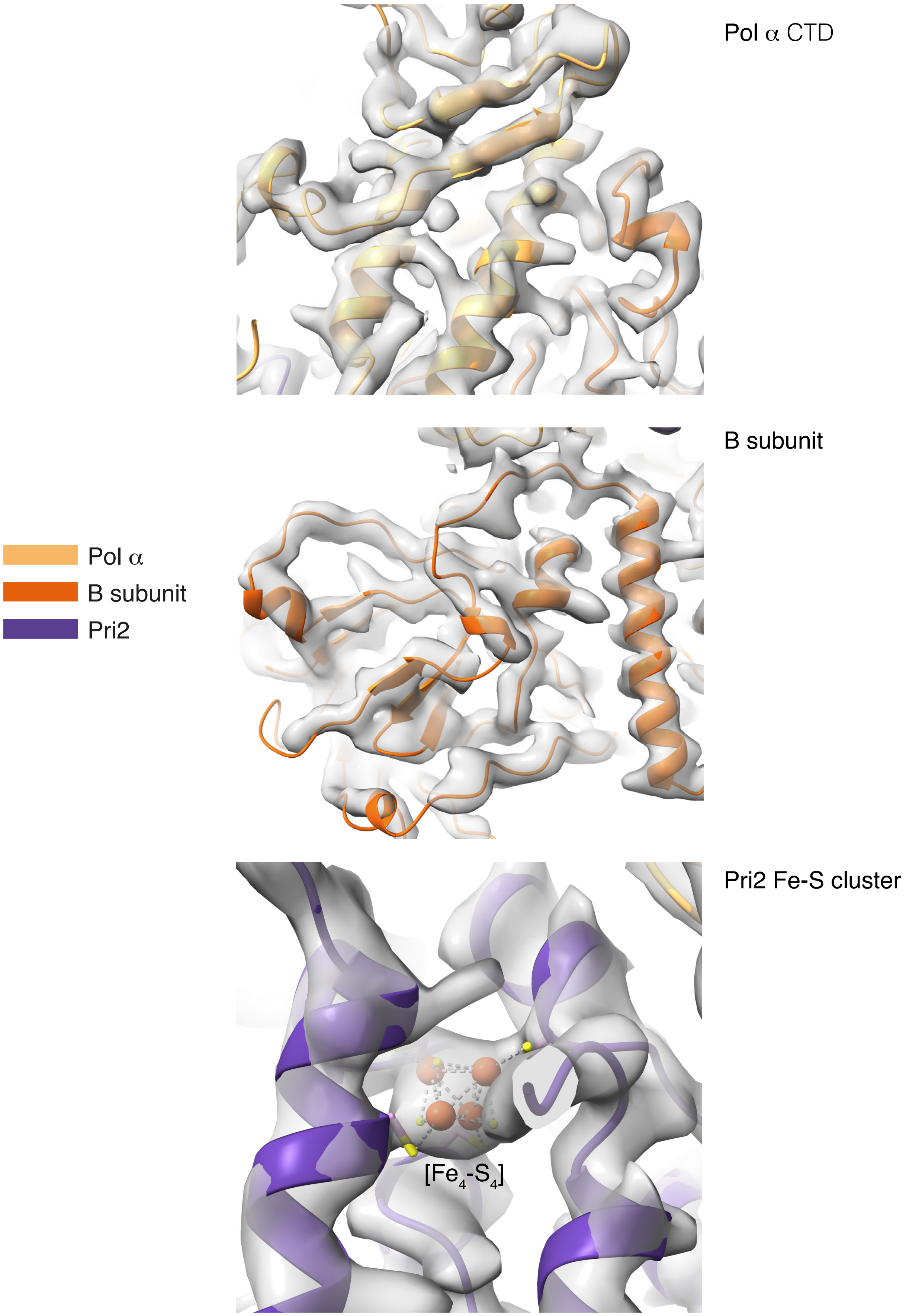

**Supplementary Figure 4**

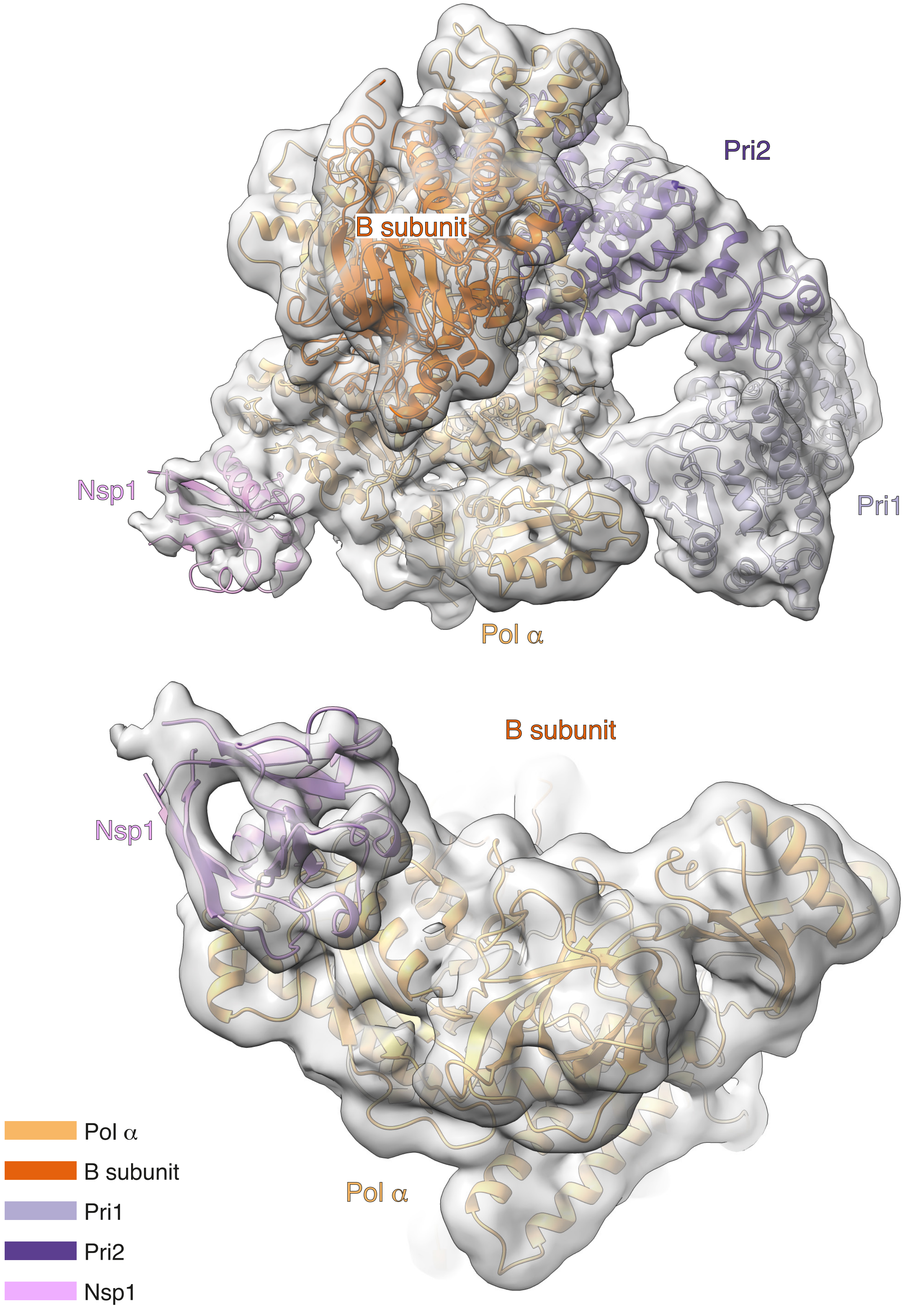

**Supplementary Figure 5**

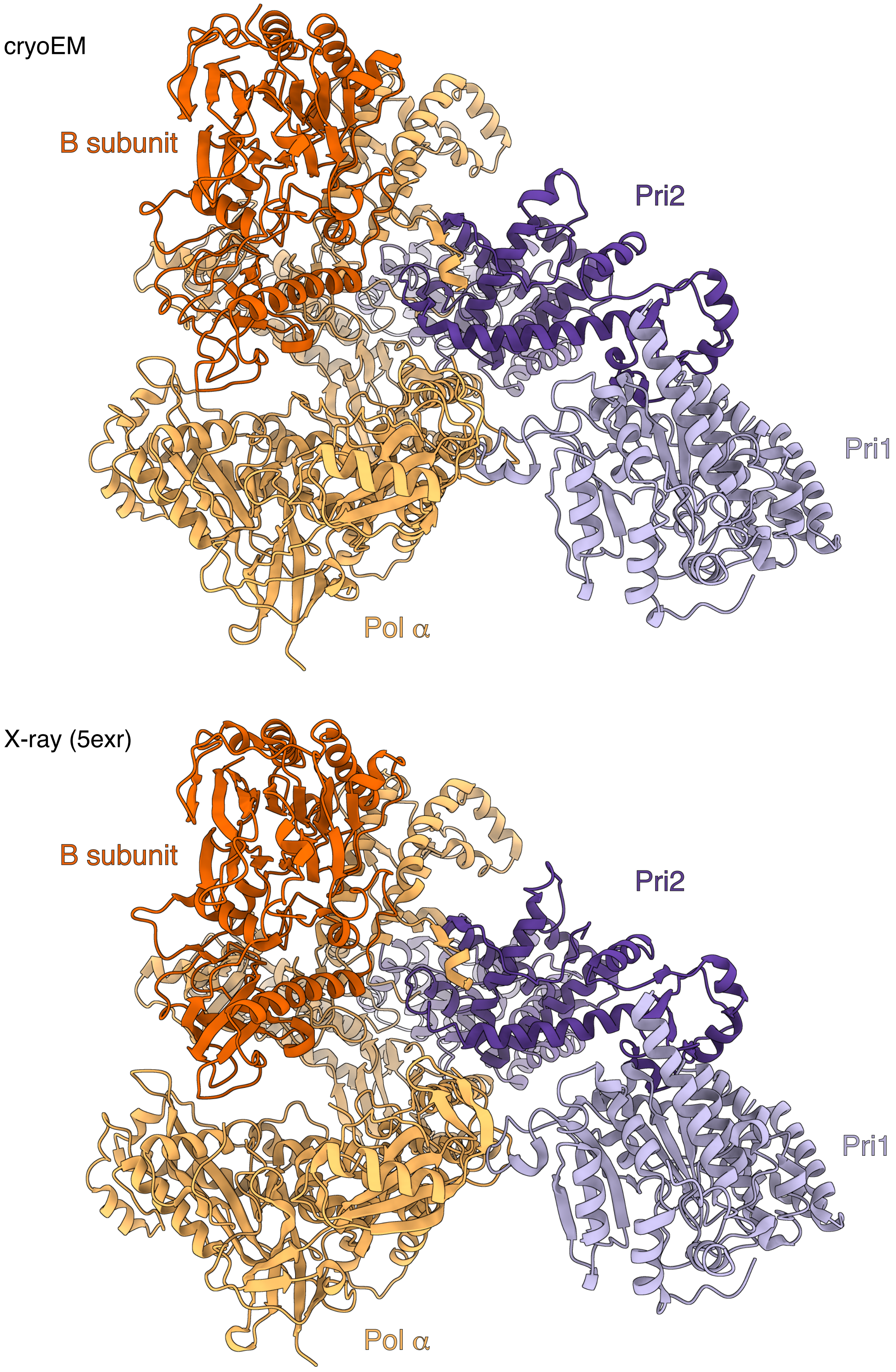

**Supplementary Figure 6**

A

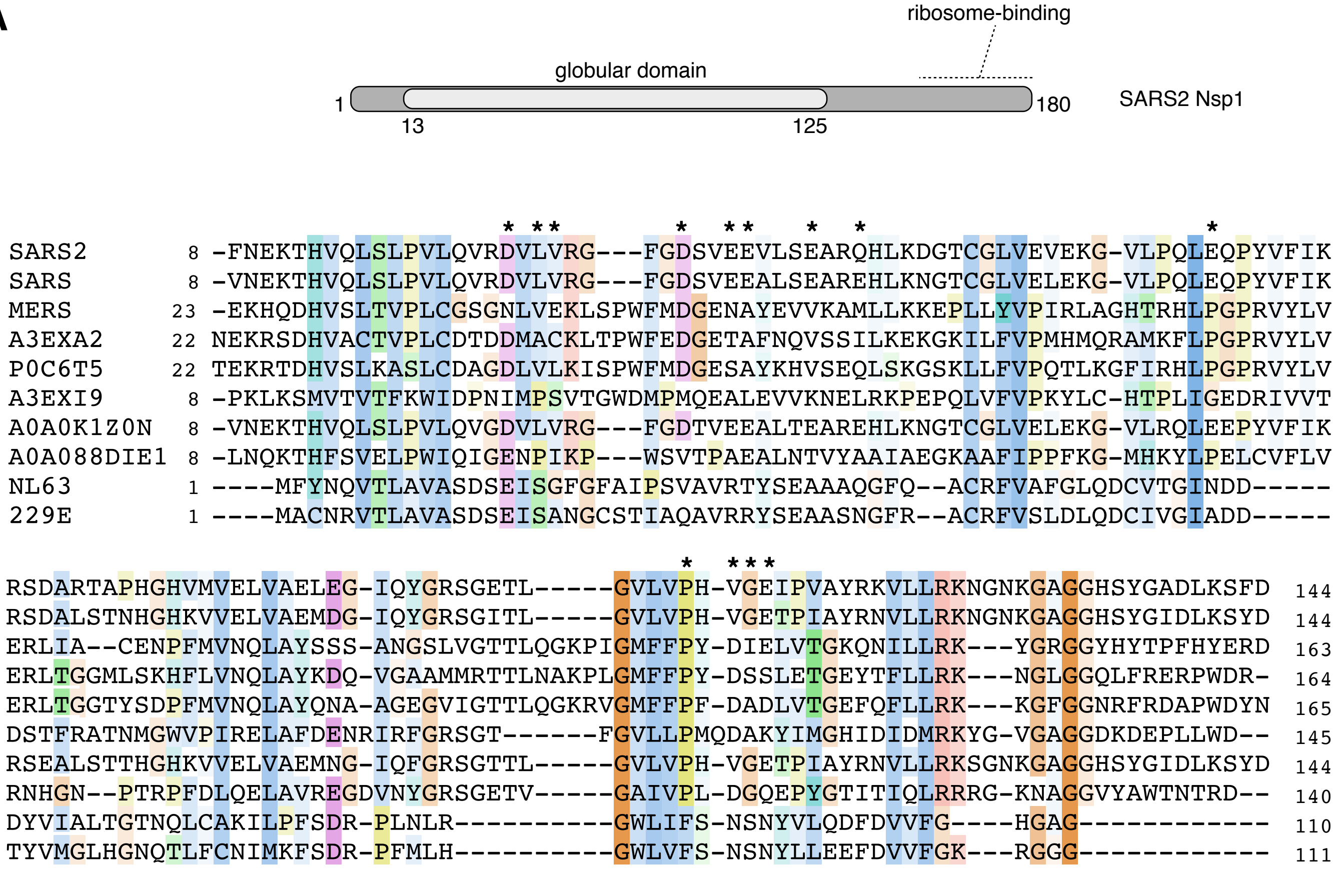

B

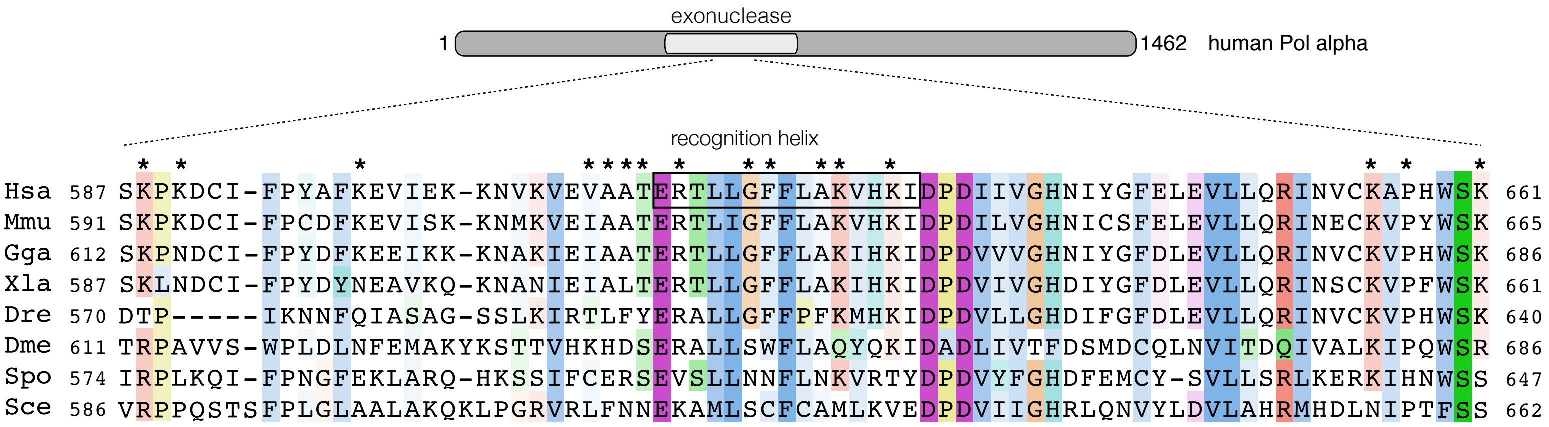

**Supplementary Figure 7**

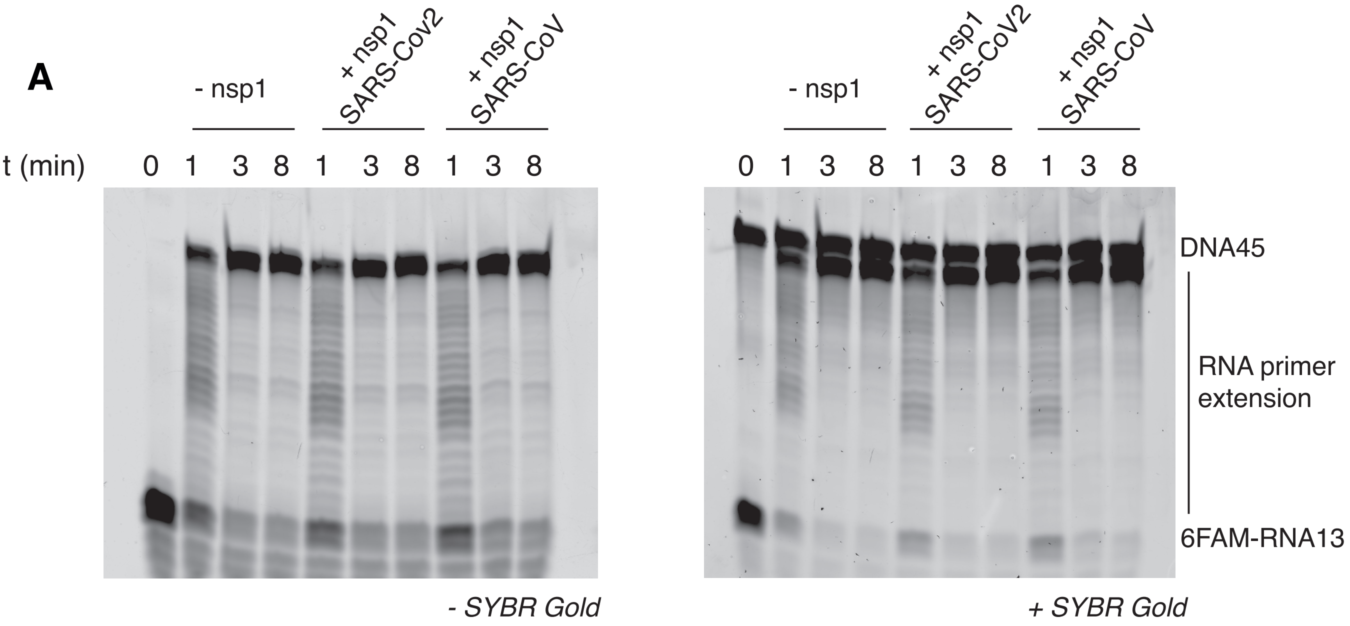

**Supplementary Figure 8**

Supplementary figure legend

**Supplementary figure 1.** Workflow of the cryoEM determination of the primosome-nsp1 structure.

**Supplementary figure 2.** Fourier shell correlation curves for masked (light grey) and unmasked (grey) 3D refined maps of the primosome - nsp1 complex and Pri1-subtracted primosome - nsp1 particles. The resolution cutoff line at 0.143 FSC is shown.

**Supplementary figure 3.** Distribution of the (rot, tilt) view angles of the primosome-nsp1 particles.

**Supplementary figure 4.** Three representative views of the sharpened map of the primosome - nsp1 complex, showing details of the Pol a CTD (top), the B subunit (middle) and the Fe-S cluster of Pri2 (purple). The primosome subunits are shown as ribbons, and the atoms as spheres.

**Supplementary figure 5.** Two views of the 3D-refined density map for the primosome - nsp1 complex, contoured at a suitable level to show the density for nsp1 and Pri1. The map in the bottom view is the 3D-refined map obtained for the Pri1-subtracted set of primosome - nsp1 particles. The structure of the complex is also shown superimposed on the density.

**Supplementary figure 6.** Comparison of the cryoEM structure (top) and X-ray crystal structure (bottom) of the human primosome.

**Supplementary figure 7.** Multiple sequence alignments of (**A**) the globular domain of related nsp1 proteins and of (**B**) the nsp1-binding sequences of the exonuclease domain of Pol α. Schematic drawings of the proteins are shown above each alignment. Interface residues involves in binding the protein partner are marked with an asterisk; the extent of the recognition helix in the nsp1-binding sequence of Pol α is indicated with a box.

A.

| | | | |
|---|---|---|---|
| SARS2 | SARS-CoV-2 | NCBI | YP_009725297.1 |
| SARS | SARS-CoV | NCBI | NP_828860.2 |
| MERS | MERS-CoV | NCBI | YP_009047213.1 |
| A3EXA2 | Bat coronavirus HKU4-2 | GenBank | ABN10847.1 |
| P0C6T5 | Bat coronavirus HKU5 | NCBI | YP_009944339.1 |
| A3EXA2 | Bat coronavirus HKU9-4 | GenBank | ABN10934.1 |
| A0A0K1Z0N | Bat SARS-like coronavirus YNLF_31C | GenBank | AKZ19074.1 |
| A0A088DIE1 | Bat Hp-betacoronavirus/Zhejiang2013 | NCBI | YP_009072438.1 |
| NL63 | Human coronavirus NL63 | GenBank | AFD98833.1 |
| 229E | Human Coronavirus 229E | GenBank | AGT21366.1 |

B.

| | | | |
|---|---|---|---|
| Hsa | Human | UniProt | P09884.2 |
| Mmu | Mouse | UniProt | P33609.2 |
| Gga | Chicken | NCBI | XP_040513999.1 |
| Xla | Frog | Uniprot | Q9DE46 |
| Dre | Zebrafish | NCBI | NP_001292393.1 |
| Dme | Fly | UniProt | P26019.2 |
| Spo | Fission yeast | UniProt | P28040 |
| Sce | Budding yeast | UniProt | P13382 |

**Supplementary figure 8.** Activity assay of the full-length human primosome in the presence of SARS or SARS2 nsp1 proteins. The two panels show the extension of a 6FAM-labelled RNA primer with dNTPs, analysed by denaturing urea acrylamide gel electrophoresis. The gel was imaged without staining (left, visualisation of 6FAM label on RNA primer) and with SYBR Gold staining (right, visualisation of all nucleic acid, including unlabelled DNA template strand).

**Supplementary Table 1. CryoEM data collection and real-space refinement**

| *Data collection* | Talos Arctica | Titan Krios |
|---|---|---|
| Voltage (keV) | 200 | 300 |
| Detector | Falcon III (counting mode) | K3 (super-resolution mode) |
| Magnification | 92,000 | 130,000 |
| Pixel size (Å/pixel) | 1.13 | 0.652 (0.326) |
| Area (Å$^2$/pixel) | 1.28 | 0.425 |
| Exposure (s) | 75 | 1.31 |
| Dose (e/pixel/sec) | 0.76 | 15.222 |
| Dose (e/Å$^2$/sec) | 0.59 | 35.81 |
| Total dose (e/ Å$^2$) | 44.31 | 46.91 |
| Number of fractions | 40 | 48 |
| Dose per fraction (e/ Å$^2$) | 1.11 | 0.98 |
| Defocus range | -2.8 -2.5 -2.2 -1.9 -1.6, -1.3, -1.0 | -2.5, -2.2, -1.9, -1.6, -1.3, -1.0, -0.7 |

| *Map resolution (Å$^2$)*[1] | Full particles | | Subtracted particles | |
|---|---|---|---|---|
| | Masked | Unmasked | Masked | Unmasked |
| $d_{99}$ | 4.07 | 4.08 | 2.94 | 2.66 |
| $d_{FSC}$ (halfmaps), 0.143 | 3.79 | 4.11 | 3.60 | 3.66 |
| $d_{model}$ | 3.90 | 3.90 | | |
| $d_{FSC\_model}$, 0.143 | 3.47 | 3.48 | | |
| $d_{FSC\_model}$, 0.5 | 4.15 | 4.16 | | |

| *Real-space refinement* | |
|---|---|
| Composition: | |
| Chains | 7 |
| Atoms | 19,748 |
| Residues | Protein: 2442 |
| Water | - |
| Ligands | SF4: 1, ZN: 3 |
| *Correlation coefficients*[1] | |
| CC, mask | 0.73 |
| CC, box | 0.79 |
| CC, peaks | 0.67 |
| CC, volume | 0.73 |
| <CC>, ligands | 0.82 |
| Bonds (rmsd): | |
| Length(Å) | 0.003 |
| Angles (°) | 0.769 |
| MolProbity score[2] | 2.07 |
| Clash score | 18.01 |

| | |
|---|---|
| Ramachandran plot (%): | |
| Outliers | 0.08 |
| Allowed | 4.46 |
| Favoured | 95.45 |
| Rotamer outliers (&) | 0.08 |
| Cβ outliers (%) | 0.00 |
| ADP (iso): | |
| Min/max/mean | |
| Protein | 63.72 / 321.38 / 153.34 |
| Nucleotide | - |
| Ligand | 122.34 / 320.81 / 145.10 |
| Water | - |

1. Afonine PV, Klaholz BP, Moriarty NW, Poon BK, Sobolev OV, Terwilliger TC, Adams PD, Urzhumtsev A. New tools for the analysis and validation of cryo-EM maps and atomic models. Acta Crystallographica Section D: Structural Biology. 2018;74(9):814–840. http://journals.iucr.org/d/issues/2018/09/00/kw5139/index.html. doi:10.1107/s2059798318009324

2. Chen VB, Arendall WB, Headd JJ, Keedy DA, Immormino RM, Kapral GJ, Murray LW, Richardson JS, Richardson DC. MolProbity: all-atom structure validation for macromolecular crystallography. Acta Crystallographica Section D: Biological Crystallography. 2010;66(Pt 1):12–21. doi:10.1107/s0907444909042073
